# Imaging of native transcription and transcriptional dynamics *in vivo* using a tagged Argonaute protein

**DOI:** 10.1101/2020.12.14.422643

**Authors:** Amel Toudji-Zouaz, Vincent Bertrand, Antoine Barrière

**Author notes:** co-last authors (Vincent Bertrand, Antoine Barrière).

## Abstract

A flexible method to image unmodified transcripts and transcription *in vivo* would be a valuable tool to understand the regulation and dynamics of transcription. Here, we present a novel approach to follow native transcription, with fluorescence microscopy, in live *C. elegans*. By using the fluorescently tagged Argonaute protein NRDE-3, programmed by exposure to defined dsRNA to bind to nascent transcripts of the gene of interest, we demonstrate transcript labelling of multiple genes, at the transcription site and in the cytoplasm. This flexible approach does not require genetic manipulation, and can be easily scaled up by relying on whole-genome dsRNA libraries. We apply this method to image the transcriptional dynamics of two transcription factors: *ttx-3* (a LHX2/9 orthologue) in embryos, and *hlh-1* (a MyoD orthologue) in larvae, respectively involved in neuronal and muscle development.

## Introduction

Understanding the dynamics of transcription is essential to deciphering cell fate specification and maintenance. While the use of fluorescent proteins as transcriptional reporters has been invaluable in this regard, it provides little information on the amount of endogenous transcripts and their dynamics. Methods such as smFISH or single-cell RNA sequencing, on fixed or dissociated tissues, allow for precise measurements of transcript abundance at the transcription site and in the nucleus, but only as a snapshot, missing information on the dynamics of transcription.

Over the last two decades, several methods were developed to label and quantify mRNA in living cells or animals (Pichon, Lagha, Mueller, & Bertrand, 2018). To this day, the most popular method for *in vivo* mRNA labelling, either of active transcription sites or individual mRNAs, is the phage-derived MS2-MCP system (E. Bertrand et al., 1998), which requires modifying the target transcript to add multiple stem-loop MS2 repeats, on which the fluorescently tagged MCP binds with high affinity. Thanks to this powerful method, over the last two decades, studies have observed dynamics and bursting transcription in cultured cells; in recent years it has been applied to live animal studies in Drosophila embryo (Bothma et al., 2014), in the mouse (H. Y. Park et al., 2014), and *C. elegans* (Lee, Shin, & Kimble, 2019). It requires, however, editing the gene of interest to insert multiple MS2 repeats or using a reporter transgene, and is difficult to scale up to assay the expression of many different genes; as a result, it has only seen limited uses in whole organisms so far. Additionally, the numerous repeats have been reported to interfere with transcript processing and cytoplasmic export (Lenstra & Larson, 2016).

More recently, alternative methods that label unmodified transcripts have been developed, first using a fluorescently labelled Pumilio homology domain (Abil, Denard, & Zhao, 2014), which can be programmed at the protein level to bind to specific 8-nucleotide RNA sequences (Ozawa, Natori, Sato, & Umezawa, 2007), then with the use of catalytically inactive Cas9 programmed with chimeric oligonucleotides (Nelles et al., 2016), or Cas13 programmed with a long guide RNA (Abudayyeh et al., 2017; L. Yang et al., 2019), to bind to a transcript of interest. These newer methods have so far only been used in cultured cells, to image accumulation of transcripts in some cellular compartments (stress granules and paraspeckles). However, to image transcription sites or single transcripts, binding of multiple fluorescent proteins on one transcript becomes necessary. Since this would require multiple guide RNAs tiling over the transcript, it renders these new methods less powerful. While efforts have expanded significantly in recent years (Pichon et al., 2018), there is still a need to develop more flexible systems to visualise native mRNAs in live cells and organisms. Particularly useful qualities would be easy programmability to bind to any RNA sequence; a good signal-to-noise ratio, to allow imaging down to single molecules; scalability up to the whole genome; and applicability to cells in culture as well as whole organisms.

Here, we report the development in the novel organism *C. elegans* of a novel method to image transcription *in vivo*, relying on the sequence-specific binding of the Argonaute NRDE-3 to transcripts in interest, that improves on existing approaches. We demonstrate its usefulness to study the transcriptional dynamics of transcription factors, in the first reported live imaging of transcription of unmodified genes in an animal. This method allows imaging both at transcription sites and in the cytoplasm, and is inexpensive to deploy.

## Methods

### Mutants

Alleles used in this study were *eri-1(mg366), nrde-2(gg95), nrde-1(gg88)*

### Transgenes

vbaIs52: _*p*_*eef-1A*.*1::YFP::nrde-3 II*

vbaIs53: _*p*_*rps-27::mNeonGreen::flag::nrde-3 II*

vbaIs54: _*p*_*eef-1A*.*1::YFP::nrde-3::SL2::sid-1 II*

vbaIs55: _*p*_*eef-1A*.*1::VenusC::nrde-3 II*

vbaIs56: _*p*_*eef-1A*.*1::VenusN::nrde-3 I*

### Plate preparation and dsRNA exposure

dsRNA targetting the gene of interest is expressed by bacteria, from the Ahringer library (Kamath et al., 2003); bacterial clones were sequenced to verify identity, and we tested the specificity to the gene of interest with Clonemapper (Thakur, Pujol, Tichit, & Ewbank, 2014). The clone targetting the gene *his-13* (ZK131.7) also matches other histone genes at several other loci across the genome.

dsRNA-expression plates were prepared fresh, at most one week before experiment. Bacteria culture expressing dsRNA against the gene of interest were plated, and the following day L3 – L4 worms added on the plate. Nuclear localisation and transcription spots were then scored on the next generation. If exposure to dsRNA persists for multiple generations, we found that transcription spots were more difficult to observe after several generations, possibly due to residual epigenetic silencing in the germline.

For certain genes, like *elt-2* or *ant-1*.*1*, primary RNAi was sufficiently efficient to affect the fitness and prevent growth. To circumvent this knockdown, we diluted the bacteria expressing dsRNA targeting the gene of interest to 50% or 75% with bacteria carrying the negative control plasmid L4440.

### Mounting and imaging

Worms were mounted on agar pads, and immobilised using levamisole or polystyrene beads. In our experience, immobilising the worms with levamisole or azide negatively affects cellular processes and hinders the measurements of transcriptional dynamics. All live imaging was performed on a spinning disk confocal microscope, with laser power between 2.5% and 16%. Image analysis was performed in Fiji; transcription spot measurements were performed in trackmate (Tinevez et al., 2017) on a maximum intensity Z projection, with background fluorescence in the nucleus being substracted from spot fluorescence.

### α-amanitin treatment

Worms were exposed to dsRNA against *hlh-1* for one generation, then L4 individuals were put in a liquid solution of α-amanitin at 20µg/mL in M9, while the negative control individuals were exposed to M9 only, for three hours before imaging.

### smFISH

Worms were grown at 20°C on dsRNA-containing NGM plates, then fixed in 4% formaldehyde/PBS for 45 minutes at room temperature, washed twice with PBS then incubated in 70% ethanol overnight at 4°C to allow permeabilisation. Embryos were equilibrated for 5 minutes with wash buffer (WB) containing 10% formamide and 2x saline-sodium citrate (SSC), then hybridised in hybridisation buffer (10% formamide, 2x SSC, 100 mg/mL dextran sulfate, 1mg/mL *E. coli* tRNA, 2 mM vanadoribosyl complex, 0.2 mg/mL BSA) containing 0.125 µM of probes, overnight at 37°C. Following hybridisation, samples were washed twice with WB including 30 minutes incubation at 37°C during the second wash then washed once in 2x SSC buffer before being mounted in Vectashield Mounting Medium with DAPI (centrifugation steps were done at 5000g). Imaging was performed by spinning disk confocal microscope.

### smiFISH

The protocol was adapted for *C. elegans* from the original publication (Tsanov et al., 2016). Briefly, mixed stages worms were fixed in 4% formaldehyde in 1x PBS for 45 minutes; rinsed twice in PBS, then permeabilised in 70% ethanol at 4°C overnight. They were then rinsed in 15% formaldehyde in 1x SSC, then incubated at room temperature for 30 minutes, before addition of the hybridisation mix (2.5µL 20x SSC, 0.8µL *S. cerevisiae* tRNA, 7.5µL formamide, 1µL probe duplexes, 13.2µL H2O; 0.5µL vanadoribosyl complex, 13.2µL 40% dextran sulfate, 10.7µL H2O) then incubated at 37°C overnight. Worms were then rinsed twice in 15% formamide in 1x SSC for 30 minutes, washed in PBS, and finally mounted in Vectashield. Spots were manually segmented in imageJ.

### qPCR

Gravid adults were bleached, and synchronized larvae were plated on dsRNA-expression plates with bacteria expressing dsRNA against *hlh-1* or GFP. Worms were harvested for RNA purification at the L4 stage using lysis reagent (Qiazol, Qiagen) according to manufacturers instructions. mRNA was reverse transcribed using SuperScript III Reverse Transcriptase (invitrogen). Quantitative PCR was carried out in CFX96 from Bio-Rad detection system with SYBR Green Master Mix (invitrogen) in 20µL total volume reaction. Primers used to quantify *hlh-1* span exons and intron boundaries, to be specific to pre mRNA, and do not match the dsRNA used. *tba-1*: TCAACACTGCCATCGCCGCC/TCCAAGCGAGACCAGGCTTCAG; *hlh-1*: ACACTGACAAGTTTCGCTGC/GAGAGCTTGAGCTTCTCCCC 3 independents biological samples were carried, each of which was measured in 3 technical replicates.

## Results

### Principle of the method

In the *C. elegans* exogenous RNAi pathway, exogenous double stranded RNA from the environment are recognized and processed into small interfering RNAs that are then loaded onto the primary Argonaute RDE-1 and bind to the target mRNA (Figure 1A). RDE-1 does not cleave the transcript (Steiner, Okihara, Hoogstrate, Sijen, & Ketting, 2009), but recruits two protein complexes: RDE-8 (Tsai et al., 2015) and RDE-12 (Shirayama, Stanney, Gu, Seth, & Mello, 2014; H. Yang et al., 2014), that degrade the target transcript, and induce recruitment of the RNA-dependent RNA polymerase RRF-1 (Aoki, Moriguchi, Yoshioka, Okawa, & Tabara, 2007). This polymerase synthesizes secondary single-stranded triphosphorylated siRNAs starting with a guanine, known as 22G siRNAs, complementary to the target mRNA, that tile over the transcript length, starting 5’ of the primary siRNA trigger (Pak & Fire, 2007; Sijen, Steiner, Thijssen, & Plasterk, 2007). These 22G siRNAs are loaded into a nematode-specific family of secondary Argonautes (Gu et al., 2009; Yigit et al., 2006) known as WAGO, mostly expressed in the germline, with the exception of the nuclear Argonaute NRDE-3 (Guang et al., 2008), expressed in the soma. Once NRDE-3 loads a 22G siRNA, it translocates from the cytoplasm to the nucleus, where it binds to the nascent transcript and recruits NRDE-2 (Guang et al., 2010), leading to blocking of transcript elongation, as well as chromatin modification through recruitment of NRDE-1 and NRDE-4 (Burkhart et al., 2011).

**Figure 1.**
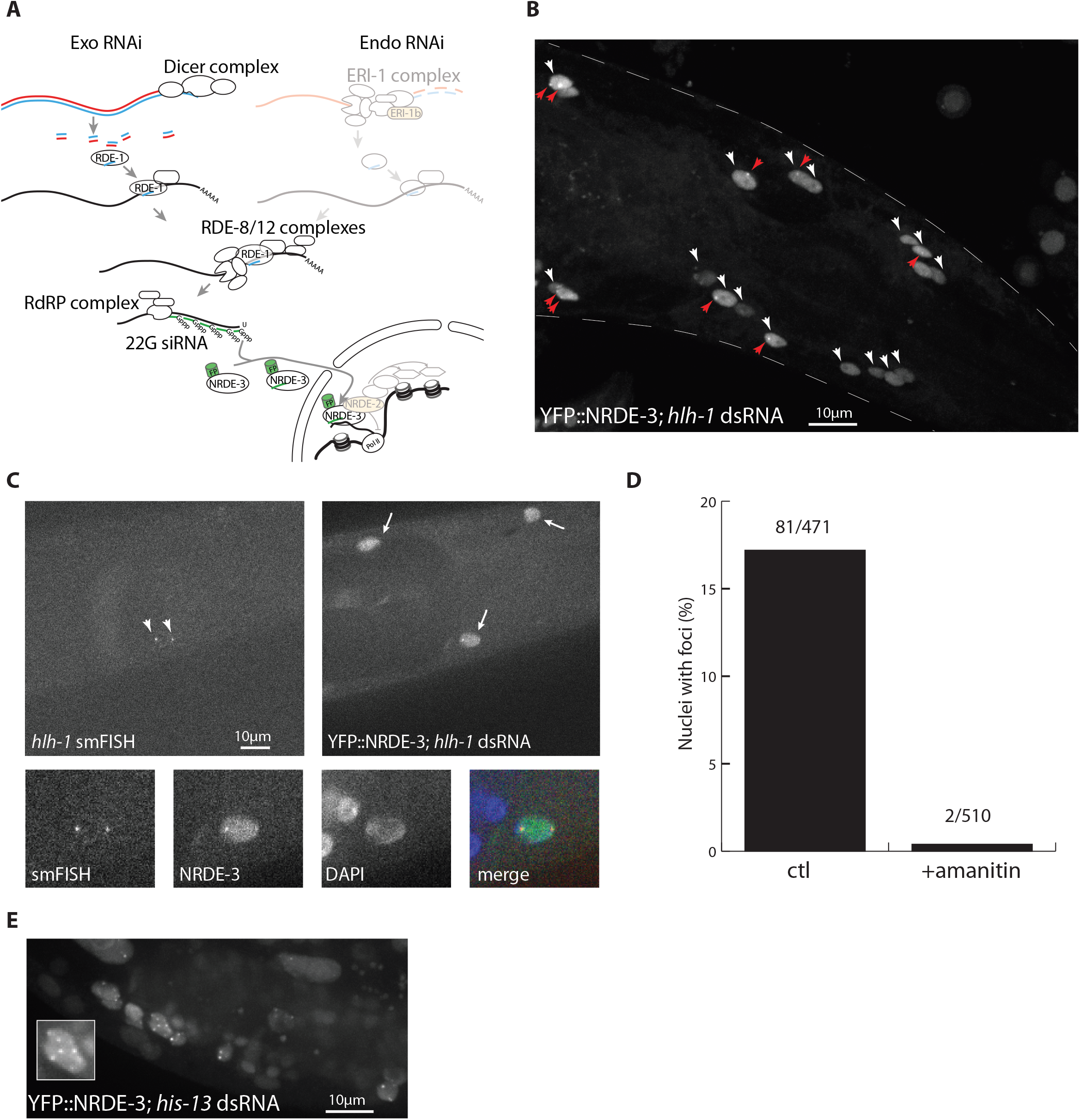
Fluorescent NRDE-3 labels active transcription sites. A. Simplified *C. elegans* RNAi pathway, as modified in our method. The exo-RNAi pathway is unaffected; the endo-RNAi pathway (upper right; greyed out) is abrogated by a mutation in ERI-1b (orange); and transcriptional silencing by the nuclear RNAi pathway (bottom right, greyed out) is blocked by a mutation in NRDE-2 (orange). NRDE-3 is labelled with a fluorescent protein (FP). B. Labelling of *hlh-1* transcription sites in larval head muscles (L4 stage). Muscle nuclei (white arrowheads) show nuclear localisation of YFP::NRDE-3, while other tissues show cytoplasmic localisation. In some of these nuclei, one or two bright foci (red arrowheads) are visible. C. Colocalisation of smFISH signal and YFP::NRDE-3. SmFISH probes matching *hlh-1* exonic and intronic sequences localise at the transcription sites. YFP::NRDE-3 localises to the nucleus of muscle cells, and accumulates at foci coinciding with smFISH spots. D. YFP::NRDE-3 foci are sensitive to amanitin. After exposure to α-amanitin, YFP::NRDE-3 foci were no longer observable. E. dsRNA matching multicopy genes induce localisation of YFP::NRDE-3 at multiple foci. dsRNA against the sequence of *his-13* also matches the sequence of other histones genes, at multiple clusters across the genome. Worms fed with dsRNA matching multiple genes displayed more than two and up to eight YFP::NRDE-3 foci, in multiple tissues (somatic gonad, hypodermis). Insert: one developing somatic gonad nucleus with seven foci.

To label active transcription sites, we took advantage of this specific binding of NRDE-3 to nascent transcripts. We introduced two mutations (Figure 1A, orange): *nrde-2(gg95)*, a putative null allele to prevent the blocking of transcript elongation (Guang et al., 2010); and *eri-1(mg366)*, to block the endogenous RNAi pathway that competes with the exogenous RNAi pathway for 22G siRNA synthesis (Guang et al., 2008). These mutations should not interfere extensively with biological processes: *eri-1(-)* is commonly used in *C. elegans* to enhance RNAi efficiency, while *nrde-2(-)* has been described as reducing brood size (Guang et al., 2010) and inducing germline mortality at 25°C (Buckley et al., 2012), but otherwise not associated with gross defects. NRDE-3 was tagged at the N-terminal end with a fluorescent protein (YFP or NeonGreen). After exposing worms to double-stranded RNA matching the sequence of a target transcript, and in the absence of transcriptional silencing due to the *nrde-2(-)* mutation, we expect to observe nuclear localisation of fluorescently-labelled NRDE-3, and accumulation of NRDE-3 at transcription sites.

This method takes advantage of several specificities of *C. elegans* RNAi pathways: the dsRNA uptake pathway (through the SID-2 (Mcewan, Weisman, & Hunter, 2012) and SID-1 (Winston, Moldowitch, & Hunter, 2002) dsRNA transporters), which allows easy delivery of dsRNA against any gene of interest in the entire animal; the RdRP complex, which synthesizes a set of 22G siRNAs antisense to the transcript of interest, 5’ from the siRNA site; existence of the cleavage-deficient nuclear Argonaute NRDE-3, that can accumulate on the nascent transcript; and the availability of genome-wide dsRNA-synthesising bacteria libraries (Kamath et al., 2003), through which worms can be exposed to dsRNA against the gene of interest simply by feeding.

### Validation of the method

In the absence of endogenous RNAi due to *eri-1(-)*, we observed cytoplasmic localisation of NRDE-3 in the majority of tissues, as previously described (Guang et al., 2008). In the germline, early embryo and intestine, however, we observed nuclear localisation in the absence of exogenous dsRNA, revealing the presence of an alternate source of 22G siRNAs in these tissues (Supplementary figure 1). When worms were exposed to dsRNA matching the sequence of the muscle-specific transcription factor *hlh-1*, the *C. elegans* orthologue of MyoD, by feeding, we observed nuclear localisation of NRDE-3 specifically in muscle cells (Figure 1B; white arrowheads), as previously described (Guang et al., 2008). In addition, while in an *eri-1(-)* background, nuclear localisation of NRDE-3 spread to all cells after a few days (Guang et al., 2008), we did not observe such generalized nuclear localisation in the *eri-1(-); nrde-2(-)* background. In a subset of muscle cells, we observed one or two bright spots in the nucleus, which we hypothesised to correspond to active *hlh-1* transcription sites (Figure 1B, red arrowheads). To demonstrate that these nuclear spots are indeed active transcription sites, we performed fluorescent *in situ* hybridization targeting *hlh-1* intronic and exonic sequences, and observed colocalisation with YFP::NRDE-3 (Figure 1C; arrowheads), thus validating the principle of our method. To show that these foci were dependent on transcription, we tested the disappearance of foci after exposure to an RNA polymerase II inhibitor. In worms exposed to the Pol II inhibitor α-amanitin (Lindell, Weinberg, Morris, Roeder, & Rutter, 1970), 2/510 nuclei showed foci, while in untreated worms, we observed foci in 81/471 nuclei (Figure 1D). Finally, when worms were exposed to dsRNA matching the sequence of highly conserved histones genes at multiple locations across the genome (ZK131.7, *his-13*), we observed more than two, and up to eight spots per nucleus (Figure 1E), as expected from the transcription site labelling of multiple histone genes. The nuclear foci therefore reflect active transcription sites.

This labelling method, to be useful, should minimally interfere with the processes it aims to image. One possible concern is interference from the tagged NRDE-3 on transcription of the gene of interest. To verify the absence of transcriptional silencing in the *nrde-2(-)* genetic background, we performed a qRT PCR on the *hlh-1* pre-mRNA, and found no evidence of downregulation in presence of dsRNA targeting *hlh-1* (Supplementary figure 2). Our results agree with previous observations showing little or no transcriptional silencing in an *nrde-2(-)* background (Guang et al., 2010). Additionally, we do not expect NRDE-3-based labelling to block splicing, since siRNAs are synthesized in the cytoplasm from mature transcripts and should not match the intron-exon boundaries present in a pre-mRNA. Finally, while Argonaute proteins do bind with high affinity to their targets, this should not be sufficient to block translation, as the similarly strong binding of MCP to the MS2 stem-loop is insufficient to block the progression of ribosomes (Halstead, Wilbertz, Wippich, & Ephrussi, 2015). A mutant for NRDE-1, which is required for transcriptional silencing downstream of NRDE-2 (Burkhart et al., 2011), instead of a *nrde-2(-)* mutation, also allowed labelling of active transcription sites (Supplementary figure 3).

We tested the applicability of this method on multiple genes, expressed at various levels in multiple tissues, and saw nuclear localisation patterns and foci corroborating independently reported expression patterns (Figure 2): wnt ligand *mom-2* in posterior cells in the mid embryo, matching previous smFISH data (Harterink et al., 2011); transcription factor *hlh-1* in muscle cells in the late embryo, matching published data (Nair, Walton, Murray, & Raj, 2013); the highly expressed adenosine nuclear transporter *ant-1*.*1* in all tissues; EGF ligand *lin-3* in the anchor cell, matching previous data (Barkoulas, van Zon, Milloz, van Oudenaarden, & Félix, 2013), GATA transcription factor *elt-2* in the intestine (Raj, van den Bogaard, Rifkin, van Oudenaarden, & Tyagi, 2008); LIM homeobox transcription factor *ttx-3* in AIY neurons in the embryo, matching previous data (V. Bertrand & Hobert, 2009); (F. Soulavie and V. Bertrand, unpublished data). The method worked in most tissues; we found that we were generally able to observe nuclear localisation, and in some of these nuclei we observed foci corresponding to active transcription sites (Supplementary table 1). This method did not work, however, with every dsRNA tested, and we failed to observe specific nuclear localisation or transcription foci for several genes, such as the notch-regulated gene *sygl-1* in the germline (Lee et al., 2019). This likely reflects the variability of RNAi efficiency, depending on tissues and dsRNA sequence.

**Figure 2.**
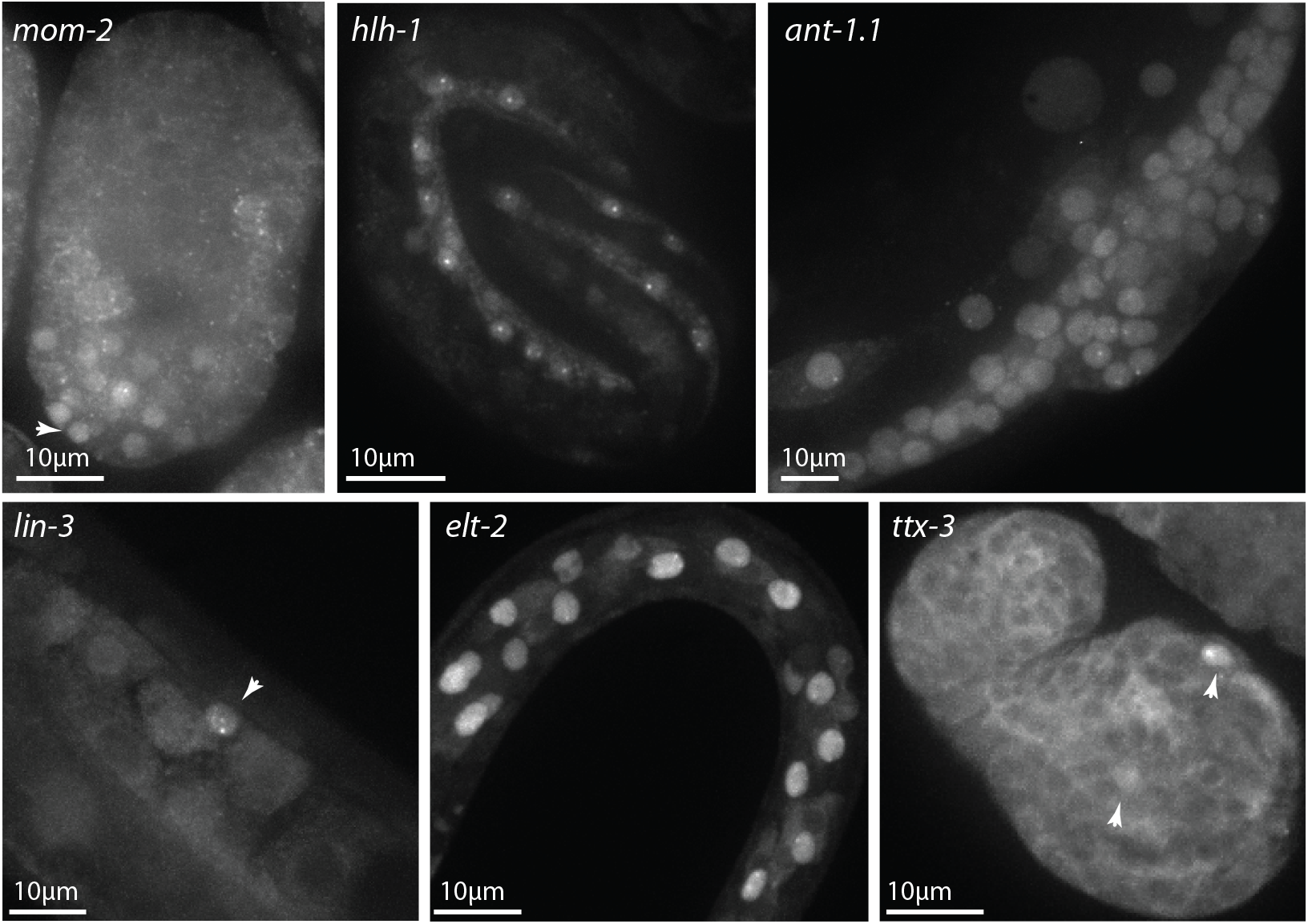
NRDE-3 labelling of transcription sites works on multiple genes and tissues. After maternal exposure to *mom-2* dsRNA, YFP::NRDE-3 is nuclear localised in the posterior part (white arrowhead) of the mid embryo, and accumulates at nuclear foci in some of these cells. After maternal exposure to *hlh-1* dsRNA, YFP::NRDE-3 is nuclear localised in muscle cells in the three-fold embryo, and accumulates at one or two transcription sites per nucleus. After exposure to *ant-1*.*1* dsRNA, all cells in this L4 larva show nuclear localisation of YFP::NRDE-3. In some of them (hypodermis, developing vulva, somatic gonad) transcription foci are visible. After exposure to *lin-3* dsRNA, the anchor cell at the third larval stage shows nuclear localisation (white arrowhead) and two transcription foci. After exposure to *elt-2* dsRNA, nuclear localisation is stronger in the intestine cells at the second larval stage, and some of them show transcription foci. After maternal exposure to *ttx-3* dsRNA, YFP::NRDE-3 is nuclear localised in AIY neurons in the comma stage embryo (white arrowheads) and, in one of them, a transcription site is visible.

### Optimisation of the method

Not all tissues were found to be equally amenable to transcriptional labelling. When a strain expressing YFP::NRDE-3 was exposed to dsRNA matching the sequence of the fluorescent protein, we would expect every fluorescent cell to have a strong nuclear signal; however, we observed that in neurons and pharynx, YFP::NRDE-3 still localised to the cytoplasm (Figure 3A). These tissues are known to have low RNAi efficiency, as they do not express the dsRNA transporter *sid-1* (Winston et al., 2002). Since expressing *sid-1* in these tissues could be sufficient to restore RNAi silencing by feeding (Calixto, Chelur, Topalidou, Chen, & Chalfie, 2010), we co-expressed *YFP::nrde-3* and *sid-1* under the control of the same *eef-1A*.*1* promoter and observed improved nuclear localisation and labelling of transcription sites in neurons and the pharynx (Figure 3B). In addition, in the case of the dsRNA against *ttx-3*, which already worked in neurons without *sid-1* expression (Figure 2), we observed a stronger nuclear localisation and clearer transcription spots (Figure 3C). Nonetheless, the general efficiency of labelling in neurons and pharynx still appeared to be lower than in other somatic tissues.

**Figure 3.**
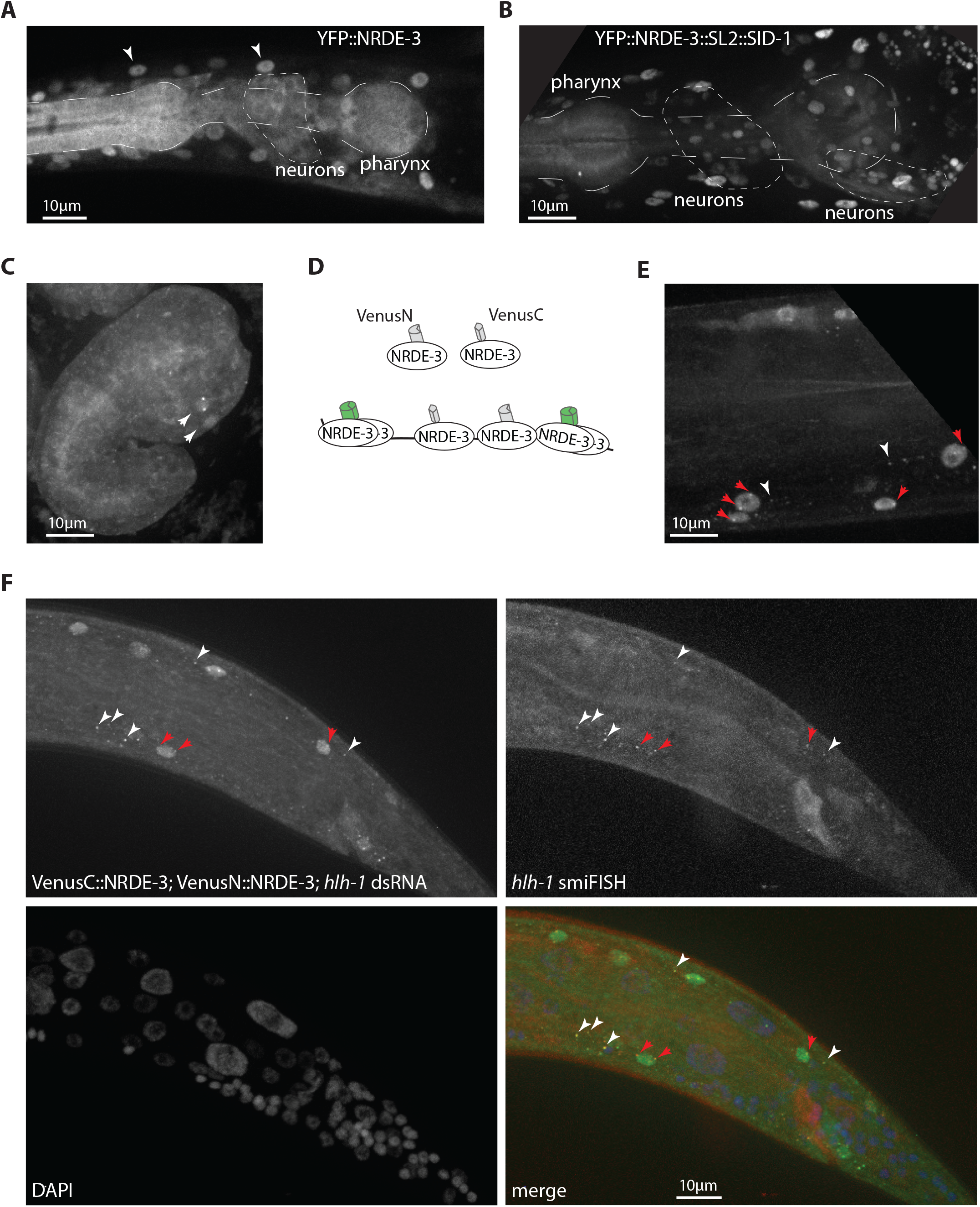
Improvements on transcriptional labelling. A. Neurons and pharynx have less effective RNAi. After exposure to dsRNA against GFP, YFP::NRDE-3 is nuclear localised in most tissues (white arrowheads: muscle nuclei), but localises to the cytoplasm in pharynx and neurons. B. Restoring expression of SID-1 in all tissues improves nuclear localisation. After exposure to dsRNA against GFP, YFP::NRDE-3 is nuclear localised in neurons and, partially, in the posterior pharynx. C. Improved transcriptional labelling in neurons. An embryo expressing SID-1 in all tissues, after maternal exposure to dsRNA against *ttx-3*, shows nuclear localisation and transcriptional foci in AIY neurons (compare with Figure 2). D. Principle of trimolecular fluorescence complementation. N-terminal and C-terminal fragments of the fluorescent protein Venus are fused to NRDE-3. In the cytoplasm or nucleus, local concentration of NRDE-3 molecules does not allow fluorescence complementation, thus reducing background fluorescence; once bound on the target transcript, VenusN::NRDE-3 and VenusC::NRDE-3 are in sufficient proximity to allow for fluorescence complementation. E. Improved labelling of transcription with trimolecular fluorescence complementation. After exposure to dsRNA against *hlh-1*, bright transcription foci are visible in multiple nuclei (red arrows); additionally, some cytoplasmic foci are visible in the cytoplasm of muscle cells (white arrowheads), but not in other tissues. F. Trimolecular fluorescence complementation labels cytoplasmic transcripts. Cytoplasmic spots (white arrowheads), due to fluorescence complementation between VenusN::NRDE-3 and VenusC::NRDE-3, as well as nuclear active transcription sites (red arrows), colocalise with smiFISH probes labelling *hlh-1* transcripts.

To improve on the method, we focused mostly on the transcription factor *hlh-1* for several reasons: first, it is transcribed throughout the life of the animal, for initiation and maintenance of the myocyte fate (Harfe, Branda, Krause, Stern, & Fire, 1998). Second, its expression starts relatively early in development, at least as early as 210 minutes (Nair et al., 2013; Packer et al., 2019), allowing for labelling in the embryo as well as larval stages. Third, as a transcription factor, its expression level is relatively low (Cao et al., 2017), providing a good test case to evaluate the applicability of our method to moderately expressed *C. elegans* genes.

While accumulation of fluorescently labelled NRDE-3 in the nucleus is useful to identify cells expressing the gene of interest, proteins not bound to a transcript have the drawback of increasing the background fluorescence above which the fluorescent signal at transcription sites needs to rise to be detectable. It is an important variable to control in transcriptional imaging: too much of the tagged protein, the transcription site or individual mRNAs cannot be seen over background fluorescence; too little, the availability of the protein may be a limiting factor (Ferguson & Larson, 2013). While sufficient for snapshot imaging, the *YFP::nrde-3* transgene we used had the drawback of high expression under control of the *eef-1A*.*1* promoter, and low photostability, restricting laser power and exposure time, thus limiting the use of time-lapse imaging. To improve these points, we tested a transgene with the more photostable and brighter fluorescent protein mNeonGreen (Heppert et al., 2016) fused to NRDE-3, under control of the ubiquitous, medium-level expression *rps-27* promoter. With the improved signal-to-noise ratio, we were able to observe more frequently transcriptional foci in several tissues (data not shown).

Another approach to reduce background fluorescence is trimolecular fluorescence complementation (TriFC), which was successfully used to label transcription sites or transcripts with other RNA binding proteins (Ozawa et al., 2007; S. Y. Park, Moon, & Park, 2020; Valencia-Burton, McCullough, Cantor, & Broude, 2007; Wu, Chen, & Singer, 2014; Yamada, Yoshimura, Inaguma, & Ozawa, 2011). We therefore decided to implement it for the NRDE-3 system. Briefly, the fluorescent protein Venus was split in two halves: VenusC and VenusN, which were fused to the N-terminal end of NRDE-3. Only when bound in close proximity to the same transcript would the two halves of Venus be close enough to reconstitute a fluorescent protein (Figure 3D). Without exposure to dsRNA, we observed a low level of nonspecific fluorescence complementation in the nerve ring, intestinal nuclei, and associated with the cytoskeleton in seam cells, but not in other larval or adult tissues. After exposure to *hlh-1* dsRNA, we observed fluorescence complementation in muscle nuclei, and bright foci in many of these nuclei (Figure 3E), with better contrast and higher frequency than observed with the simpler *YFP::nrde-3* transgene. Occasionally, we also observed additional fainter nuclear spots, possibly representing mRNAs being processed for export, as well as cytoplasmic spots, which we hypothesized to be cytoplasmic transcripts (Figure 3E). We therefore performed smiFISH labelling (Tsanov et al., 2016) targeting *hlh-1*, and observed colocalisation in both nuclei and cytoplasm (Figure 3F), confirming their nature as cytoplasmic transcripts. We observed co-labelling by NRDE-3 and smiFISH probes for 82% of cytoplasmic transcripts; a ratio similar to those observed with double labelling with two smFISH probe sets targetting the same transcript (Raj et al., 2008; Stapel et al., 2016; Tsanov et al., 2016), indicating efficient and specific labelling of single mRNAs with NRDE-3. Additionally, we did not observe qualitative changes of transcript localisation as a consequence to NRDE-3 based labelling (Supplementary figure 4). Reconstituted Venus is however less photostable than the other fluorescent proteins, and therefore less suitable for timelapse imaging.

### Use of the method to monitor transcriptional dynamics of transcription factors

We next used this method to follow the dynamics of transcription over time in living animals. We first exposed nematodes expressing YFP::NRDE-3 and SID-1 in all tissues to dsRNA against the transcription factor *ttx-3*, an LHX-2/9 orthologue necessary for acquisition of the AIY neuronal fate, and imaged their progeny at intervals of four minutes, to minimise bleaching. We observed, in embryos, nuclear localisation of the fluorescent signal in AIY neurons and followed the appearance and disappearance of transcription spots during embryonic development (Figure 4A). By tracking the intensity over time, we were able to measure transcriptional activity (Figure 4B) and observe, in one embryo, a maximum of activity around 56 minutes after spot appearance, thereafter decreasing and disappearing 32 minutes later. We next analysed the dynamics of *hlh-1* transcription factor, to evaluate the possibility of tracking transcription in multiple nuclei simultaneously. To improve the signal-to-noise ratio, reduce phototoxicity and bleaching, we used a strain expressing mNeonGreen::NRDE-3. We imaged worms at the L4 stage for two hours and recorded the intensity of nuclear foci over time. We were able to follow the appearance and disappearance of transcriptional foci, and quantify changes in intensity (Figure 4C). We observed that, within one worm, transcriptional foci in body wall muscle nuclei appeared within 20 minutes of each other, and appear to be correlated. We observed oscillation of apparent transcription levels, with a high transcription state ranging from 18 to 32 minutes. There were also brief troughs in transcription or complete disappearance of fluorescent foci (Figure 4C, nucleus 2) of transcription spots. In nuclei where we observed two transcription spots (Figure 4C, nucleus 3), we saw no correlation in intensity fluctuation between the two loci. The data indicate that our method allows precise monitoring of transcriptional dynamics.

**Figure 4.**
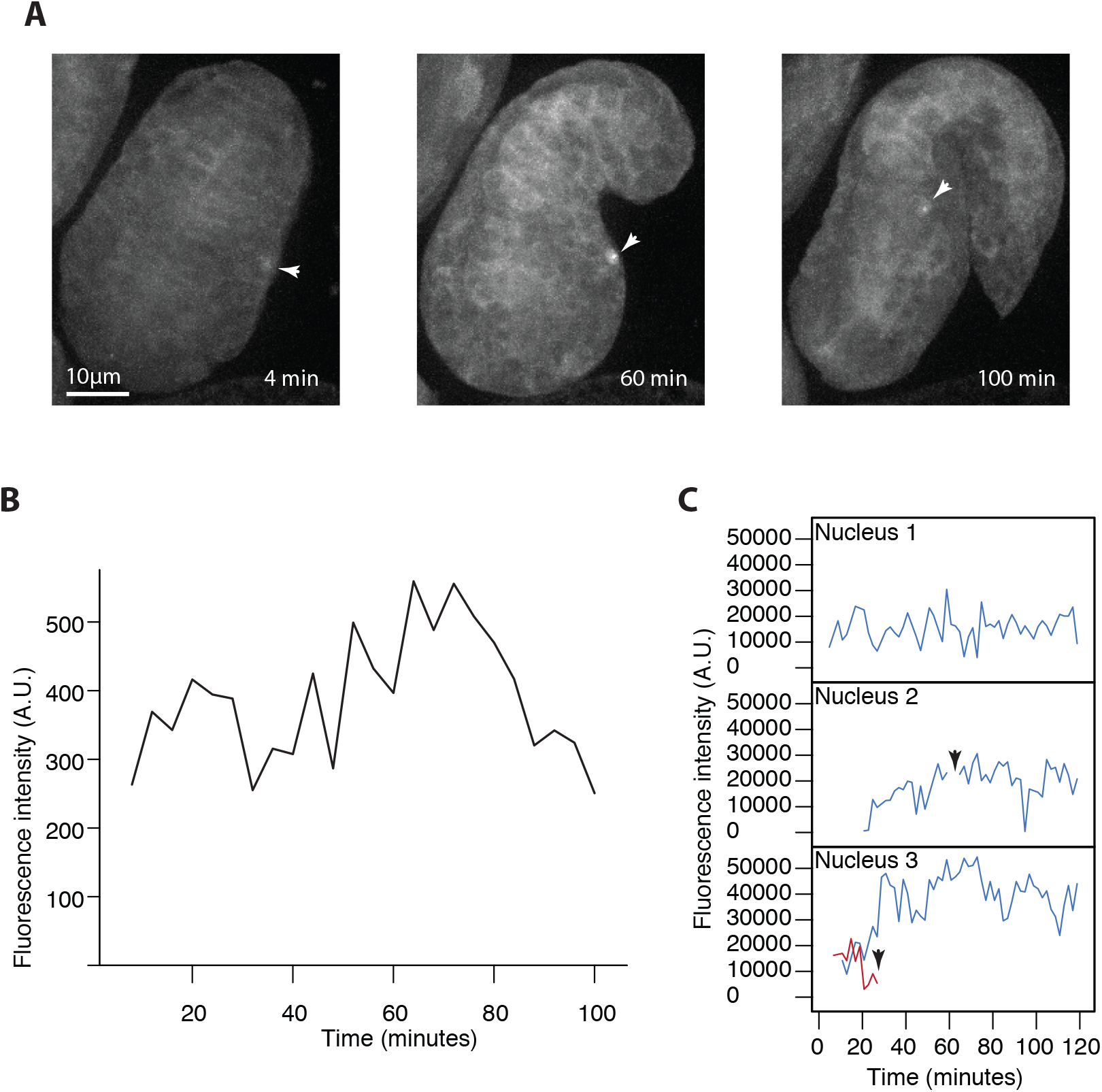
NRDE-3 labelling reveals dynamics of transcription. A. Dynamics of *ttx-3* transcription in the embryo. Three frames showing the AIY nucleus (white arrowheads) before a transcription spot is visible, at 4 minutes; a bright transcription spot at 60 minutes, and a dimming spot, before it dropped below detection limit, at 100 minutes. B. Fluorescence intensity of the spot seen in panel A graphed over time. The spot is visible starting at t=8 minutes, reaches maximum intensity at 64 minutes, and drops below detection level after 100 minutes. Spot detection threshold by eye is around 200 A.U. C. Fluorescence intensity of *hlh-1* transcription spots in three nuclei. Some transcription spots decrease at or around the background fluorescence level, and seem to disappear (nuclei 2 and 3; arrowheads). In nucleus 3, two transcription spots were tracked concurrently until one disappeared.

## Discussion

Here, we developed and applied a novel method to image transcription in *C. elegans*. This is, to our knowledge, the first dynamic imaging of transcription, without modification of the gene of interest, in a live animal. We took advantage of the *C. elegans* RNAi pathways and tools to program NRDE-3 to bind to the transcript of interest. This approach makes it a fast and inexpensive method, bypassing the need for *in vitro* synthesis of antisense RNAs and their delivery into cells of interest. Our approach offers several advantages over current *in vivo* transcription imaging methods. Contrary to the popular MS2 method, it does not require modification of the transcript of interest, which can perturb the system and cannot be easily scaled up. In addition, our method induces binding of multiple fluorescent proteins to each target transcript from a single dsRNA trigger, allowing detection of low quantities, down to single mRNA molecules. This result would be difficult to obtain with methods relying on programmable proteins such as Pumilio, Cas9 or Cas13, which would require the introduction of either multiple programmed Pumilio transgenes or long RNAs.

Although powerful, this method has several identified limitations. Because synthesis of 22G siRNAs is downstream of primary RNAi, which requires transcript cleavage by RDE-8/12 complexes to recruit the RdRP complex (Figure 1A), some level of post-transcriptional silencing can still be present. To mitigate this silencing of target genes when necessary, we reduced the amount of dsRNA ingested by diluting the dsRNA-expressing bacteria of interest with bacteria expressing a dsRNA not matching any *C. elegans* sequence. Additionally, because the target gene needs to be already expressed for synthesis of 22G siRNAs to proceed, there exists a delay between the first initiation of transcription of the gene of interest and translocation of fluorescently labelled NRDE-3 to the nucleus, allowing for labelling of transcription sites.

The expression patterns we observed with NRDE-3 labelling are fully consistent with data independently obtained by other teams. Comparisons with smFISH or smiFISH labelling of *hlh-1* transcripts, the most accurate mRNA visualisation methods, indicate that NRDE-3 labelling accurately represents endogenous expression and transcript localisation (Figures 1C, 3F; supplementary figure 4), with similar efficiency. In the absence of a preexisting method to follow transcriptional dynamics of unmodified genes in live animals, it is more difficult to establish directly validating comparisons for the transcriptional dynamics we observed. Nonetheless, transcriptional dynamics of other genes observed in whole animals with the MS2-MCP reporter transgene system revealed transcriptional bursts of the notch target gene *sygl-1* in the *C. elegans* germline, ranging from 20 to 70 minutes (Lee et al., 2019); in the early *D. melanogaster* embryo, MS2-MCP imaging of the *eve* stripe 2 enhancer revealed short transcription bursts of 4 to 10 minutes (Bothma et al., 2014). The dynamics here reported for transcription factors *ttx-3* and *hlh-1* are in a range compatible with those observed for other genes involved in fate determination.

Our method will allow us, among other things, to study the dynamics of terminal transcription factors responsible for cell fate acquisition, and how their bursty transcription affects fate specification and maintenance. The apparent correlated appearance of *hlh-1* transcription foci in different nuclei we observed could indicate that *hlh-1* may be under the control of the heterochronic pathway, which regulates the precise timing of gene expression during larval stages.

While we developed this novel method for use in *C. elegans*, it could be transposed to other organisms with adjustments to account for divergence of RNAi pathways. Beyond its use as an imaging platform, we demonstrated the usefulness of the Argonaute NRDE-3 as a general purpose, highly specific, RNA programmable RNA binding protein, which opens multiple possible applications for transcriptome engineering (Mackay, Font, & Segal, 2011). Argonautes bind with high affinity and specificity to their target mRNAs (Wee, Flores-Jasso, Salomon, & Zamore, 2012), making it a tool of choice to be developed further. While previous attempts at protein complementation with Argonaute proteins *in vitro* did not give specific results (Furman et al., 2010), the success of trimolecular fluorescence complementation on transcripts shows that NRDE-3 could also be used for split protein complementation *in vivo*, for example to activate proteins only in target cells or at specific genomic loci (Shekhawat & Ghosh, 2011; Wei & Wang, 2015).

## Acknowledgments

We thank the CGC for strains and reagents; J. Ewbank, E. Bertrand and X. Pichon for extensive discussions and suggestions; members of our lab for comments on the manuscript, and F. Soulavie for communicating unpublished results. Imaging was performed on PiCSL-FBI core facility (IBDM, AMU-Marseille) supported by the French National Research Agency through the “Investments for the Future” program (France-BioImaging, ANR-10-INBS-04). Some strains were provided by the CGC, which is funded by NIH Office of Research Infrastructure Programs (P40 OD010440). This work is funded by the Agence Nationale de la Recherche (ANR-11-LABX-0054 and ANR-17-ERC2-0018) and the Fondation pour la Recherche Médicale (DEQ20180339160).

---

**Supplementary figure 1.**
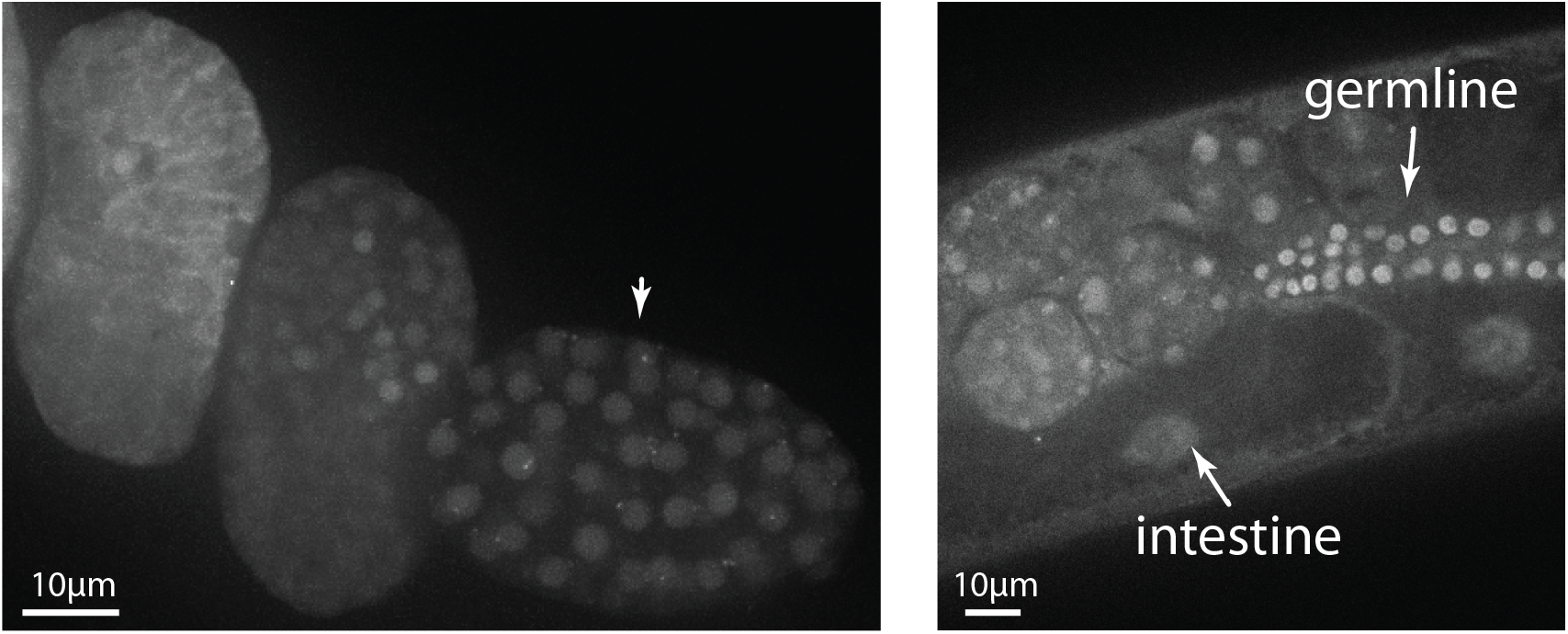
Nuclear localisation of YFP::NRDE-3 in intestine, germline, and early embryo in the absence of dsRNA exposure. Nuclear localisation in a *nrde-2(-); eri-1(-)* background in the early embryo until around 100 cell stage (left), intestine and germline in adults (right), in the absence of dsRNA exposure. Note presence of perinuclear spots in the early embryo (arrowheads).

**Supplementary figure 2.**
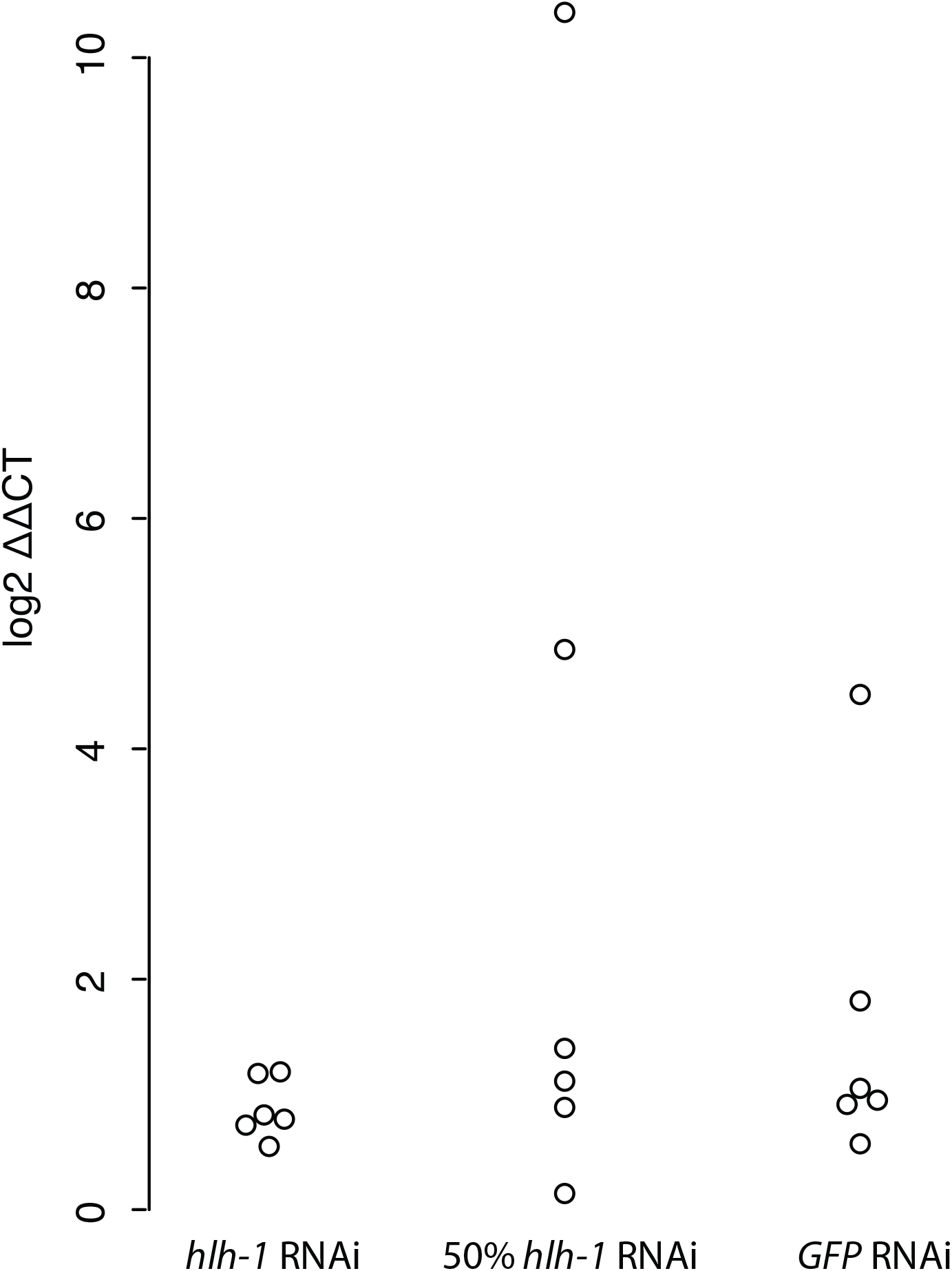
Absence of significant *hlh-1* downregulation in the *nrde-2(-)* background. After exposure to dsRNA against GFP (as control), against *hlh-1* or against a mixture of the two, qRT-PCR failed to find differences in *hlh-1* transcription level relative to *tba-1*. T-test between the control *GFP* RNAi, and *hlh-1* and 50% dilution *hlh-1* RNAi found no statistically significant difference (p=0.26 and 0.41, respectively).

**Supplementary figure 3.**
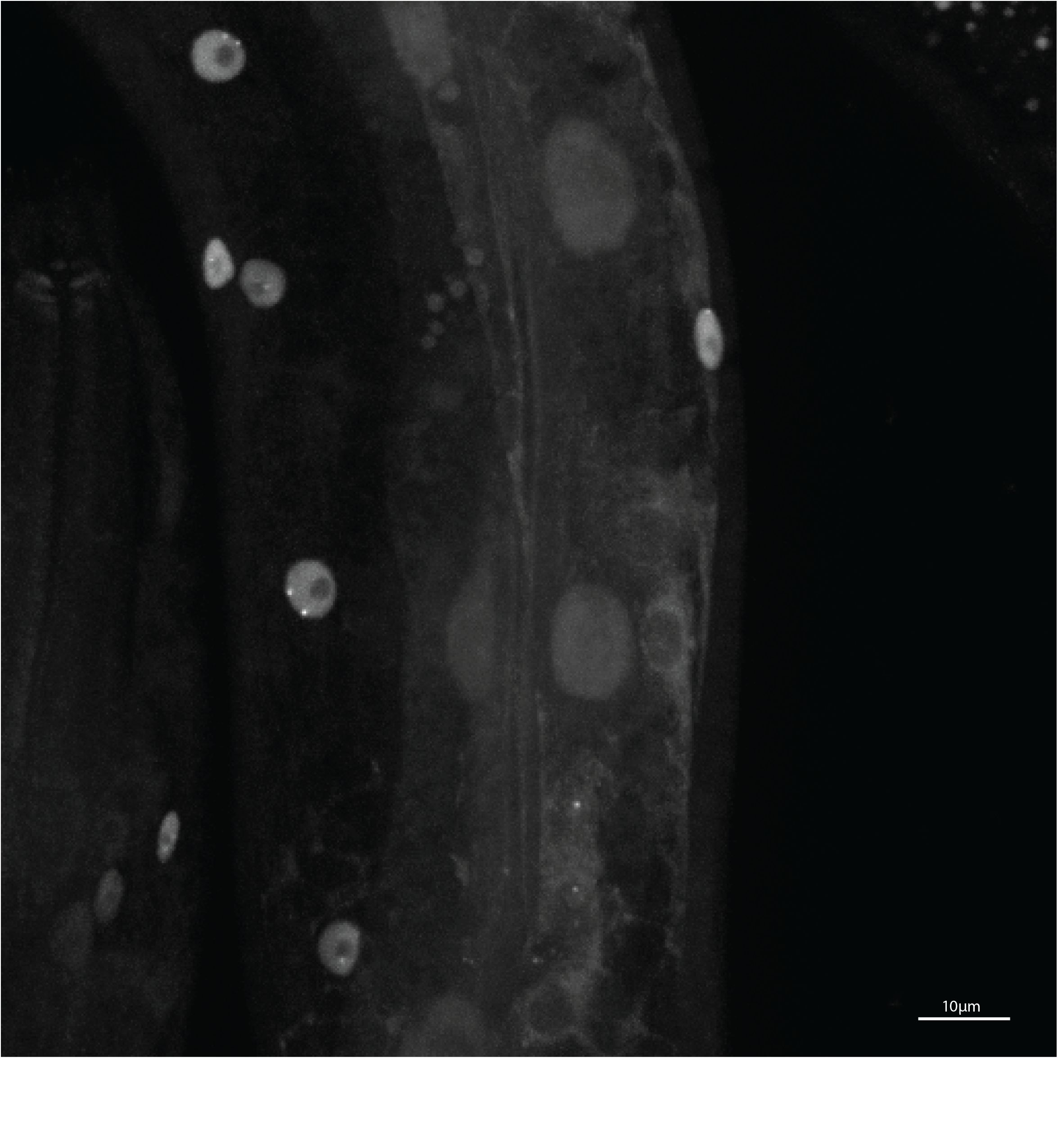
*nrde-1* mutation also allows labelling of transcription sites. After exposure to *hlh-1* dsRNA, we observed nuclear localisation, and labelling of transcription sites, in nematodes of the genotype *eri-1(-); nrde-1(-); yfp::nrde-3*.

**Supplementary figure 4.**
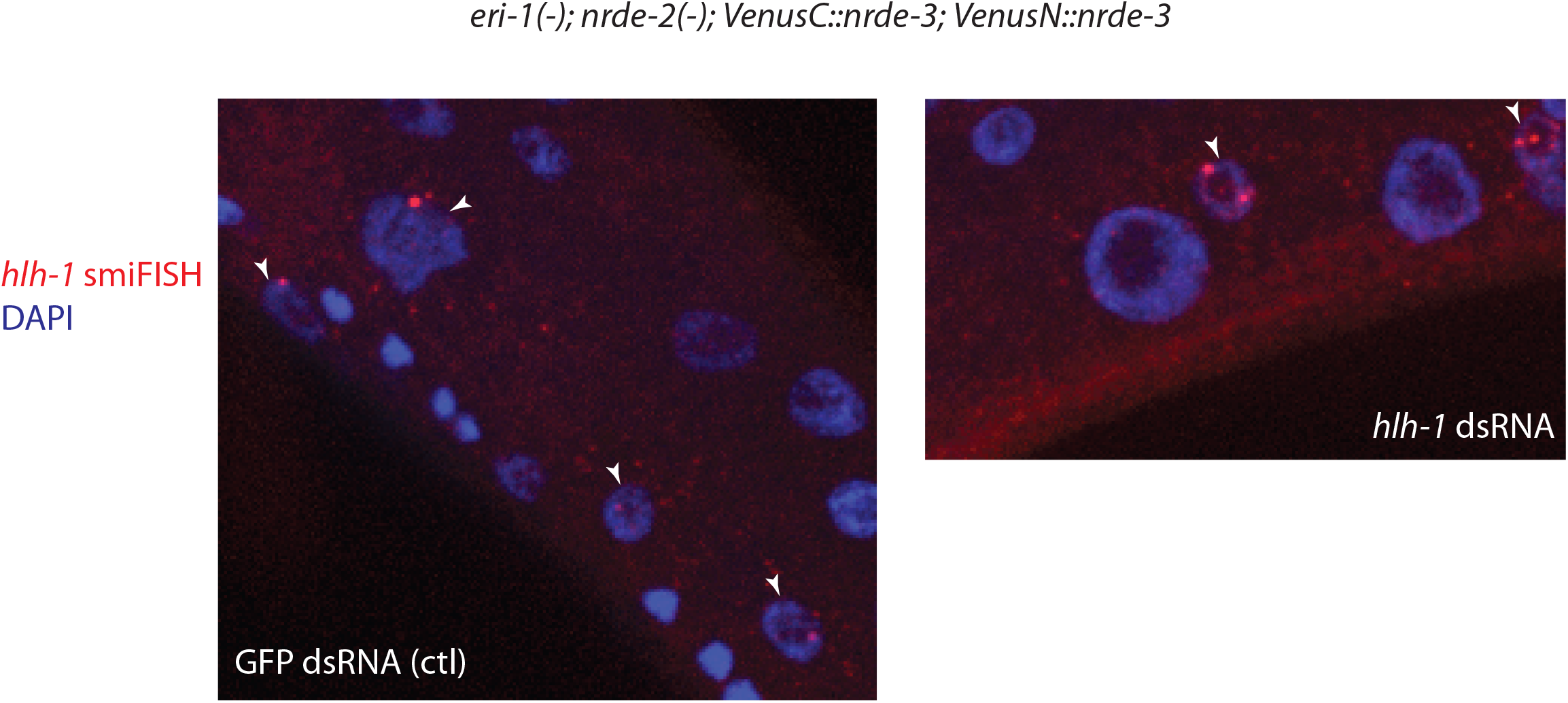
NRDE-3-based labelling does not qualitatively affect transcript localisation. After exposure to dsRNA against *hlh-1* or to a control dsRNA (against GFP), *hlh-1* transcript localisation is not qualitatively different; after smiFISH labelling (red), we observed similar bright transcription foci in muscle nuclei (white arrowheads), and single transcripts throughout the cytoplasm. Nuclei are labelled with DAPI (blue).

**Supplementary table 1. Genes for which we observed nuclear localisation or nuclear foci after exposure to dsRNA.**

