## Supplementary material for "Imaging of native transcription and transcriptional dynamics *in vivo* using a tagged Argonaute protein": Table S1

| Gene | Expression pattern | Observed |
| --- | --- | --- |
| *ahr-1* (C41G7.5) | Multiple neurons | Nuclear localisation, no foci observed |
| *ama-1* (F36A4.7) | All cells | Nuclear localisation and foci |
| *ant-1.1* (T27E9.1) | All cells | Nuclear localisation and foci |
| *ceh-10* (W03A3.1) | Multiple neurons | Nuclear localisation, no foci observed |
| *che-1* (C55B7.12) | ASE neurons | Nuclear localisation and foci |
| *dpy-7* (F46C8.6) | Hypodermis | Nuclear localisation and foci |
| *eef-1A.1* (F31E3.5) | All cells | Nuclear localisation and foci |
| *elt-2* (C33D3.1) | Intestine | Nuclear localisation and foci |
| *his-13* (ZK131.7) | All cells | Nuclear localisation and foci |
| *his-30* (F35H10.1) | All cells | Nuclear localisation and foci |
| *his-8* (F45F2.12) | All cells | Nuclear localisation, no foci observed |
| *his-9* (ZK131.3) | All cells | Nuclear localisation and foci |
| *hlh-1* (B0304.1) | Myocytes | Nuclear localisation and foci |
| *lag-2* (Y73C8B.4) | DTC | Nuclear localisation and foci |
| *lin-3* (F36H1.4) | Anchor cell | Nuclear localisation and foci |
| *mom-2* (F38E1.7) | Posterior early embryo | Nuclear localisation and foci |
| *nlp-29* (B0213.4) | Hypodermis | Nuclear localisation and foci |
| *sygl-1* (T27F6.4) | Distal germline | No nuclear localisation |
| *unc-30* (B0564.10) | Multiple ventral cord neurons | No nuclear localisation |

Supplementary table 1
